# The co-repressor Groucho limits progression through the early transcription elongation checkpoint in vivo

**DOI:** 10.1101/2022.09.17.508372

**Authors:** María Lorena Martínez Quiles, Barbara H Jennings

## Abstract

Promoter-proximal pausing of RNA polymerase II (RNAP II) at the early elongation checkpoint is a key regulatory step in developmental and stimulus-responsive gene expression. How pausing is established and modulated in a gene-specific manner during animal development remains unclear. The Groucho (Gro)/Transducin-like Enhancer of split (TLE) family of co-repressors is widely used by transcription factors to repress transcription, yet the mechanism of Gro-mediated repression in vivo is unresolved. Here, we combine genome-wide chromatin profiling with in vivo genetic analysis in *Drosophila melanogaster* to test whether Gro regulates RNAP II pausing. Analysis of ChIP-seq data across distinct cell types shows that Gro recruitment is largely cell-type specific but consistently occurs as discrete peaks within accessible, enhancer-associated chromatin. Gro occupancy frequently overlaps promoters enriched for pausing regulators, including Negative Elongation Factor (NELF), GAGA Factor (GAF), and components of Positive Transcription Elongation Factor b (P-TEFb), without excluding their recruitment. Using a sensitised wing-specific knockdown assay, we demonstrate that partial depletion of NELF subunits, GAF, and 7SK snRNP components synergistically enhances *gro* phenotypes beyond additive effects. These genetic interactions support a shared role in regulating transcription during development. Our findings support a model in which Gro attenuates transcription by modulating progression through the P-TEFb-dependent early elongation checkpoint in vivo.

## INTRODUCTION

The precise control of gene expression in response to developmental and environmental cues is essential for the survival of all organisms. This regulation occurs primarily at the level of transcription, where the DNA template is transcribed into messenger RNA (mRNA) by RNA polymerase II (RNAP II). Transcription factors (TFs) bind to regulatory elements within the DNA to either activate or repress RNAP II activity. While transcriptional activators have been studied extensively, the mechanisms by which TFs function as repressors remain less understood. *Drosophila* has provided a powerful model for uncovering fundamental principles of transcriptional regulation during development. Although certain epigenetic features differ from mammals, the core machinery and regulatory strategies controlling RNA polymerase II transcription are widely conserved (Core and Adelman, 2019; Small and Arnosti, 2020). During development, the expression of key genes involved in cell fate determination is often governed by the spatially restricted localization or activity of repressive TFs. Notably, many of these factors lack intrinsic repressive activity; instead, they recruit co-factors (co-repressors) to inhibit productive transcription (Ferrie *et al*., 2022).

The Groucho (Gro) family of co-repressors is recruited by diverse transcription factor families during animal development (Jennings and Ish-Horowicz, 2008; Agarwal, Kumar and Mathew, 2015). This family is highly conserved across eukaryotic evolution, with functional homologues identified in organisms ranging from plants and yeast to mammals. While *Drosophila* possesses a single orthologue (*gro*), the human and mouse genomes contain four paralogous genes (*TLE1-4* and *Grg1-4*, respectively), representing a functional expansion of the family in vertebrates. The recruitment of Groucho (Gro) and its mammalian TLE orthologues is mediated by a diverse profile of transcription factors (TFs), including members of the Hes, Runx, Nkx, LEF1/Tcf, Pax, Six, and Fox families. These interactions predominantly occur via short linear amino acid motifs: the internal engrailed homology-1 (eh1) motif (FxIxxIL), and the WRPW/Y tetrapeptides. Both these motifs have been shown to bind the surface of the central pore of the β-propellor formed by the WD-domain of Gro (Jennings *et al*., 2006; Jennings and Ish-Horowicz, 2008). Systematic characterisation of the *Drosophila* proteome identified eh1 motifs in 48 TFs (Klaus *et al*., 2022). In addition, there are approximately 20 *Drosophila* TFs that contain the WRPW/Y motifs (including members of the Hes and Runx families), and TFs known to interact with Gro via other sequences including dTCF (Mieszczanek, Roche and Bienz, 2008) and Capicua (Cic) (Jiménez *et al*., 2000). These recruitment motifs and interactions with Gro family proteins are conserved across mammalian transcription factor families, consistent with the evolutionary conservation and functional importance of Gro-mediated repression.

Gro family proteins are key effectors of signalling pathways that control cell fate specification including Notch and Wnt and participate in a diverse range of biological processes including neurogenesis, somitogenesis, osteogenesis, haematopoiesis, stem cell maintenance and cancer pathogenesis (Jennings and Ish-Horowicz, 2008; Agarwal, Kumar and Mathew, 2015). They can serve as integration points for signalling crosstalk during cell fate specification (Hasson *et al*., 2004; Hasson and Paroush, 2006). For example, in the developing *Drosophila* wing, EGFR signalling promotes vein formation, whereas Notch signalling antagonises this fate (Blair, 2007). EGFR activation triggers MAPK-dependent phosphorylation of Gro, attenuating its corepressor activity and relieving repression of vein-promoting genes. In neighbouring intervein cells, Notch-induced E(spl)-bHLH repressors and Cic recruit Gro to repress these targets and enforce intervein identity (Hasson and Paroush, 2006). More recently, Gro has been identified as a component of the cell cycle regulatory network downstream of Cdk1 activity (Bar-Cohen *et al*., 2023). During S phase, Gro represses genes including *e2f1*, preventing inappropriate acceleration of cell cycle progression. Subsequent relief of Gro-mediated repression by Cdk1 is required for proper progression through G2 and entry into mitosis.

Genome-wide profiling of Gro occupancy has provided insight into how it can confer rapid and reversible repression in response to cell signalling and cell cycle cues. Although *Drosophila* Gro functions as a co-repressor, its binding is not enriched in regions marked by classical repressive chromatin modifications. Instead, Gro is most frequently recruited to genomic regions characterised by transcription factor occupancy and chromatin modifications associated with active, transcriptionally permissive chromatin. (Kaul, Schuster and Jennings, 2014; Chambers *et al*., 2017). Gro occupancy within active chromatin suggests that it does not primarily mediate repression through orchestrating large-scale chromatin restructuring. Rather, Gro may exert its repressive effects, at least in part, through effects on activity of the RNAP II complex.

The transition from transcription initiation to elongation involves a regulatory checkpoint that constitutes a major control point for many developmentally regulated genes (Jonkers and Lis, 2015; Core and Adelman, 2019; Dollinger and Gilmour, 2021). This checkpoint can be regulated to repress or activate gene expression. When progression through this step is rate-limiting, RNAP II accumulates near the transcription start site and is described as promoter-proximally paused. The phenomenon of RNAP II pausing is widespread in *Drosophila* and mammalian cells, especially at genes involved in developmental and stimulus-responsive pathways. However, how developmental cues establish or maintain pausing in a gene-specific manner remains poorly understood (Jennings, 2013; Gaertner and Zeitlinger, 2014; Core and Adelman, 2019). While promoter swapping experiments in *Drosophila* indicated that sequences around the promoter are sufficient to invoke RNAP II pausing (Lagha *et al*., 2013), a model for how promoter elements regulate RNAP II pausing remains elusive (Dollinger and Gilmour, 2021).

Molecular data to date indicate a role for Gro in gene-specific RNAP II pausing, but further evidence is required to establish the biological relevance of this mechanism. Previously, we and others have shown that Gro is frequently recruited to genes that exhibit RNAP II pausing in *Drosophila* cell lines and embryos (Kaul, Schuster and Jennings, 2014; Chambers *et al*., 2017). Functional depletion of Gro provided evidence that it modulates progression through the early elongation checkpoint at specific target genes (Kaul, Schuster and Jennings, 2014). For example, depletion of Gro from *Drosophila* Kc167 cells does not change the overall amount of RNAP II recruited at the TSS at a known target gene *E(spl)mβ-HLH*, but it does lead to an increase in RNAP II Ser2-P (the elongating form of RNAP II) and corresponding release of RNA polymerase II pausing and increased full-length transcript production (Kaul, Schuster and Jennings, 2014).

Here, we extend our previous analysis of the genome-wide recruitment of Gro to show that recruitment to open chromatin is a general feature and while there are common Gro target sites across cell types, most Gro recruitment is cell-type specific. In addition, Gro recruitment frequently overlaps that of factors known to regulate RNAP II pausing and *gro* genetically interacts with genes encoding these factors during both wing development. Taken together, our results support a model where Gro mediates repression of target genes by inhibiting RNAP II progression through the early elongation checkpoint.

## RESULTS

### Genome-wide profile of Gro recruitment in ML-DmBG3-c2 cells

We have previously profiled the recruitment of Gro in Kc167 and S2R+ cell lines (Kaul, Schuster and Jennings, 2014). Both lines were derived from *Drosophila* embryos and have properties related to plasmatocytes (phagocytic blood cells) (Cherbas *et al*., 2011). To determine which features of Gro recruitment are shared with cells derived from a different tissue, we have extended our analysis of Gro recruitment to the ML-DmBG3-c2 (BG3) cell line. BG3 cells were derived from the central nervous system (CNS) of third instar larvae (Cherbas *et al*., 2011) and were characterized extensively for genome-wide chromatin modifications as part of the modENCODE project, enabling comparison of Gro recruitment with genome-wide maps of chromatin features and regulatory factor occupancy (Celniker *et al*., 2009; Kharchenko *et al*., 2011). Transcriptomic analyses have shown that *Drosophila* cell lines retain stable, lineage-associated gene expression programs and maintain distinct transcriptional identities, allowing comparison of independently generated datasets within and between cell types (Cherbas *et al*., 2011).

To profile genome-wide Gro binding in BG3 cells, we performed ChIP-seq using an anti-Gro antibody as described previously for Kc167 and S2R+ cells (Kaul, Schuster and Jennings, 2014). Gro binding sites were determined by combining peaks with a q value of < 0.1 that were present in two biological replicates (see materials and methods for further details). Comparison of Gro peaks in BG3, Kc167 and S2R+ cells revealed extensive cell-type–specific recruitment of Gro (Figure 1A). There were 831 (from total 1248) peaks unique to BG3 cells and only 175 peaks that were found in all three cell types. 133 of the Gro peaks in BG3 cells overlap Gro peaks in Kc167 cells but not peaks in S2R+ cells. Conversely, 109 peaks found in BG3 cells overlap with peaks in S2R+ cells but not Kc167 cells. Given the diversity of transcription factors that recruit Gro and the distinct transcription factor repertoires expressed in each cell type, the observed cell-type specific binding patterns are consistent with recruitment driven by cell-type specific transcription factor expression.

**Figure 1:**
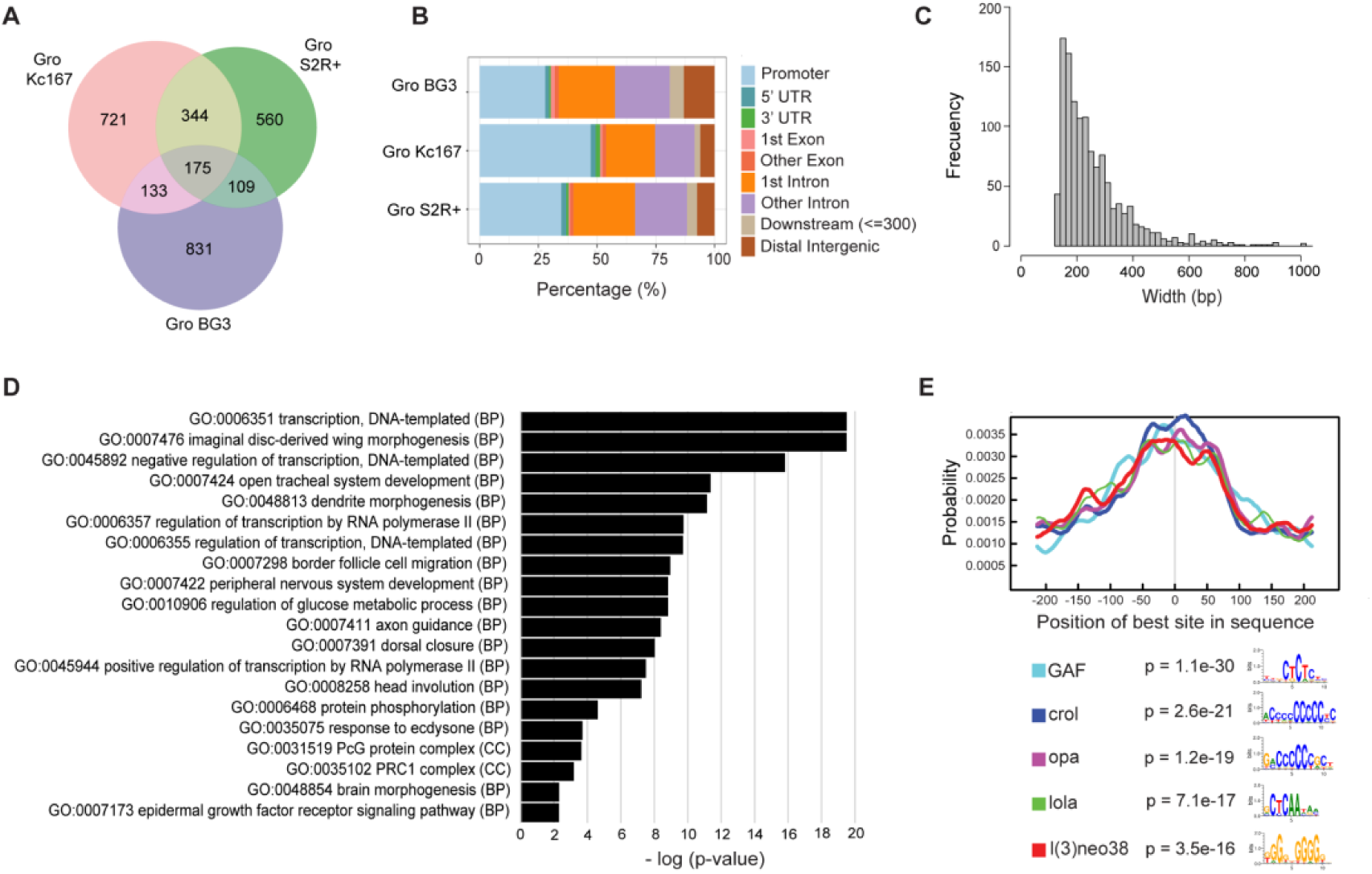
Characterization of high confidence Gro binding sites in BG3 cells and comparison among Gro peaks in three different cell lines. A) Venn diagram visualizing the overlap among Gro peaks in the Kc167, S2R+ and BG3 cell lines. Overlap was defined by a minimum and reciprocal fraction of 5% overlap across peaks. B) Bar graph showing the percentage of Gro peaks within each genomic annotation in the Kc167, S2R+ and BG3 cell lines. Promoters were defined as ±250 bp with respect to TSSs. C) Histogram showing the distribution of Gro peak widths. D) DAVID Gene Ontology (Huang, Brad T. Sherman and Lempicki, 2009; Huang, Brad T Sherman and Lempicki, 2009) analysis of genes associated with Gro binding sites in BG3 cells. Terms plotted were selected by taking the most significant term in a cluster (p-value < 10-5) and the most significant unclustered terms. E) Graph showing the top-scoring five results from the CentriMo analysis at Gro binding sites in BG3 cells: GAF, Crol, Opa, Lola, and l(3)neo38 (Bailey and Machanick, 2012). Each curve represents the probability of the binding location for the named factors at each position of Gro ChIP-seq peak regions (500 bp). The legend shows the motifs and its central enrichment p-value.

Gro is predominantly recruited to targets in a cell specific manner, yet its distribution across genomic features is similar in BG3, Kc167 and S2R+ cells (Figure 1B). In all three cell types, Gro peaks are enriched at promoter and intronic regions, consistent with the profile previously observed in embryos (Chambers *et al*., 2017). In BG3 cells, Gro binding is detected as discrete peaks rather than broad domains, in agreement with previous observations in Kc167 and S2R+ cells (Kaul, Schuster and Jennings, 2014). Gro peaks in BG3 cells have an average peak width of 278 bp and a median peak width of 224 bp (Figure 1C). In all three cell lines Gro binding is detected as discrete, focal peaks rather than extended domains, which is not consistent with the historical model in which Gro spreads across chromatin from an initial site of transcription factor-mediated recruitment (Chen and Courey, 2000). Despite differences in target loci between cell types, the positional bias and focal nature of Gro recruitment are conserved.

Gene Ontology analysis indicated that the Gro peaks in BG3 cells were associated with transcripts linked to a variety of developmental processes including imaginal disc derived wing morphogenesis, open tracheal system development, and dendrite morphogenesis, in addition to transcripts linked with transcriptional regulation (Figure 1D). Motif analysis of sequences underlying Gro peaks (Figure 1E) revealed enrichment for GAGA Factor (GAF) binding sites, as observed in Kc167 and S2R+ cells, and for l(3)neo38 sites shared with S2R+ cells (Kaul, Schuster and Jennings, 2014). Motifs for Crooked legs (Crol), Odd paired (Opa) and Longitudinals lacking (Lola) were additionally enriched in BG3 cells but were less frequent in Kc167 or S2R+ cells. These differences in sequence context are consistent with recruitment by distinct transcription factor repertoires in different cell types.

The limited overlap of Gro peaks and the distinct motif enrichments across the three cell lines indicate that Gro binding is largely cell type specific. Nevertheless, in all three cell types Gro is recruited to discrete peaks located predominantly at promoters and within intronic regions. Thus, while target identity varies with transcription factor expression, the positional bias and focal nature of Gro recruitment are conserved.

### Gro is enriched in open “Enhancer” type chromatin

To gain insight into the molecular mechanism underlying Gro-mediated repression, we analysed the chromatin environment associated with Gro peaks. Although it is well established that Gro acts as a repressor, recent studies have indicated that Gro predominantly binds in chromatin that permits transcription. The integrative analysis of the binding profiles of 53 proteins (tagged with DamID) that associate with chromatin in Kc167 cells produced a model in which the *Drosophila* genome contains five principal chromatin types; ‘‘Red’’ (active, developmentally regulated), ‘‘Yellow’’ (active, housekeeping), ‘‘Blue’’ (repressed, by Polycomb Group complexes) ‘‘Green’’ (repressed, classic heterochromatin), and ‘‘Black’’ (highly repressed) (Filion *et al*., 2010). We previously observed that Gro recruitment in Kc167 cells is enriched in Red chromatin and overlaps DNase I hypersensitive regions, consistent with recruitment to active rather than repressed chromatin (Filion *et al*., 2010; Cockerill, 2011; Kaul, Schuster and Jennings, 2014).

To determine whether recruitment to active chromatin is a general feature of Gro binding, we extended this analysis using additional data from Kc167 cells and by examining BG3 cells. Analysis of ATAC-seq data (Buenrostro *et al*., 2013) for Kc167 cells (Porcelli *et al*., 2019) confirmed our previous observation that Gro is predominantly recruited to open chromatin (Figure 2A); just 4.4% of Gro peaks did not overlap ATAC-seq peaks (Supp Figure 1). Having established that Gro is recruited to open chromatin in Kc167 cells, we extended our analysis to determine if this holds true across different cell types by analysing DNase I hypersensitivity data from BG3 cells. Akin to Kc167 cells, Gro peaks were also found in regions associated with DNase I hypersensitivity in BG3 cells (Figure 2B). Thus, in both embryonic and CNS-derived cell lines, Gro binding is associated with open chromatin rather than with chromatin states classified as repressed.

**Figure 2:**
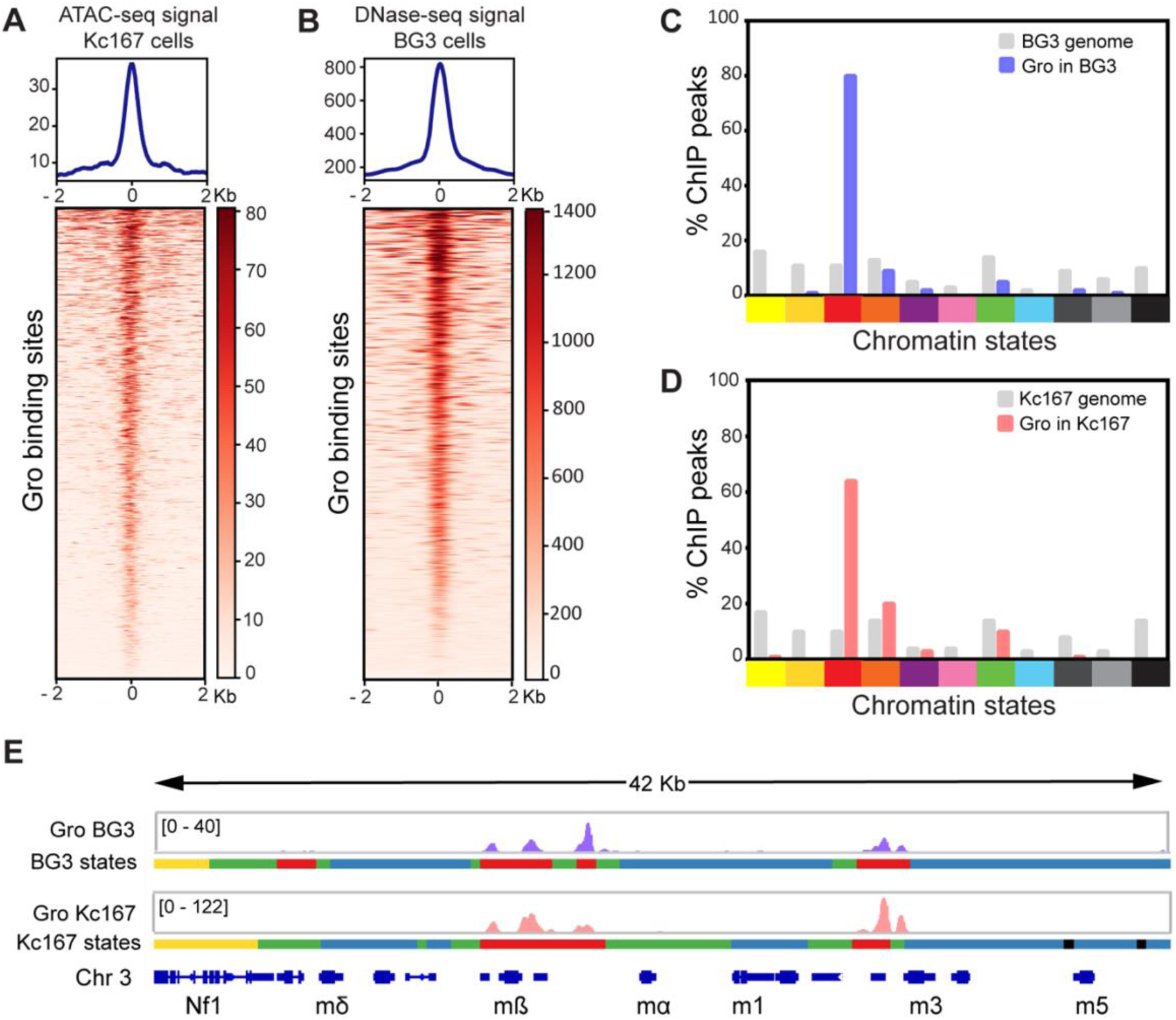
Gro binding sites are present at regions of open chromatin that are enriched in “enhancer” chromatin in Kc167 and BG3 cells. A) Average profile plot and heatmap showing the ATAC-seq signal at Gro peaks in Kc167 cells. B) Average profile plot and heatmap showing the DNase-seq signal at Gro peaks in BG3 cells. The average plots and the heatmaps are centred at the centre of Gro peaks and extended 2 Kb upstream and downstream of that reference point. C) Distribution of Gro binding sites according to their chromatin type in BG3 and D) Kc167 cells. Chromatin states are coloured according to the scheme defined by (Skalska *et al*., 2015). Gray bars indicate the proportion of each chromatin signature in the total *Drosophila* genome in each cell line. E) Gro binding profiles in BG3 and Kc167 cells along the *E(spl)-C* locus aligned with the chromatin map colour coded as defined by (Skalska *et al*., 2015).

To further define the chromatin context of Gro recruitment, we compared Gro peaks with chromatin state maps generated in Kc167 and BG3 cells (Skalska *et al*., 2015). These maps were generated from data from 24 different chromatin modifications and DNase I hypersensitive sites and led to the definition of 11 chromatin states using an adaptation of the Hidden Markov model (HMM) approach (Kharchenko *et al*., 2011). We observed that Gro peaks are most highly enriched in “Enhancer” (Enh) chromatin (red) in both BG3 and Kc167 cells (Figure 2C and D). Enh chromatin is characterised by the H3K4me1, H3K27ac and H3K56ac modifications and DNAse I hypersensitivity and is one of the three defined states of active regulatory chromatin. Around 80% of Gro peaks mapped to Enh chromatin in BG3 cells and over 60% of the superset of high confidence Gro peaks in Kc167 cells defined by (Kaul, Schuster and Jennings, 2014). Gro binds predominantly in Enh (red) chromatin at the established target genes, *E(spl)mβ-HLH* and *E(spl)m3-HLH* in both cell lines (Figure 2E) (Kaul, Schuster and Jennings, 2014). There was also some enrichment of Gro peaks in “Active TSS” (orange) in Kc167 cells, but Gro peaks were largely excluded from chromatin states associated with transcriptional repression, including Polycomb, heterochromatin and basal states (Figure 2C, D).

Taken together, all these observations indicate that Gro is primarily recruited to chromatin that is accessible and permissive for transcription, rather than chromatin that is compacted and repressed. We therefore next examined whether Gro represses transcription by modulating progression through the early elongation checkpoint rather than by altering chromatin accessibility.

### Gro is recruited to promoters enriched for regulators of RNAP II pausing

Previous studies have revealed that Gro recruitment is associated with target genes that exhibit promoter proximal RNAP II pausing and depletion of Gro can lead to pause release (Kaul, Schuster and Jennings, 2014; Chambers *et al*., 2017). These observations suggest that Gro functions in the context of promoter proximal pausing. To determine whether Gro binding sites overlap genomic binding profiles of regulators of RNAP II pausing, we compared Gro peaks with peaks of factors known to promote pausing and pause release.

Recruitment of the NELF complex is linked with RNAP II pausing at genes in *Drosophila* (Lee *et al*., 2008) and acts to promote pausing (Yamaguchi *et al*., 1999). We examined the overlap of Gro peaks with the Nelf-E peaks generated by ChIP-seq in Kc167 cells by (Nazer *et al*., 2018). We observed that 64.7% of Gro peaks overlap with Nelf-E genome-wide and that 91.2% of Gro peaks located at TSSs overlap with Nelf-E peaks (Figure 3A), indicating that Gro frequently occupies promoters where NELF is also detected.

**Figure 3:**
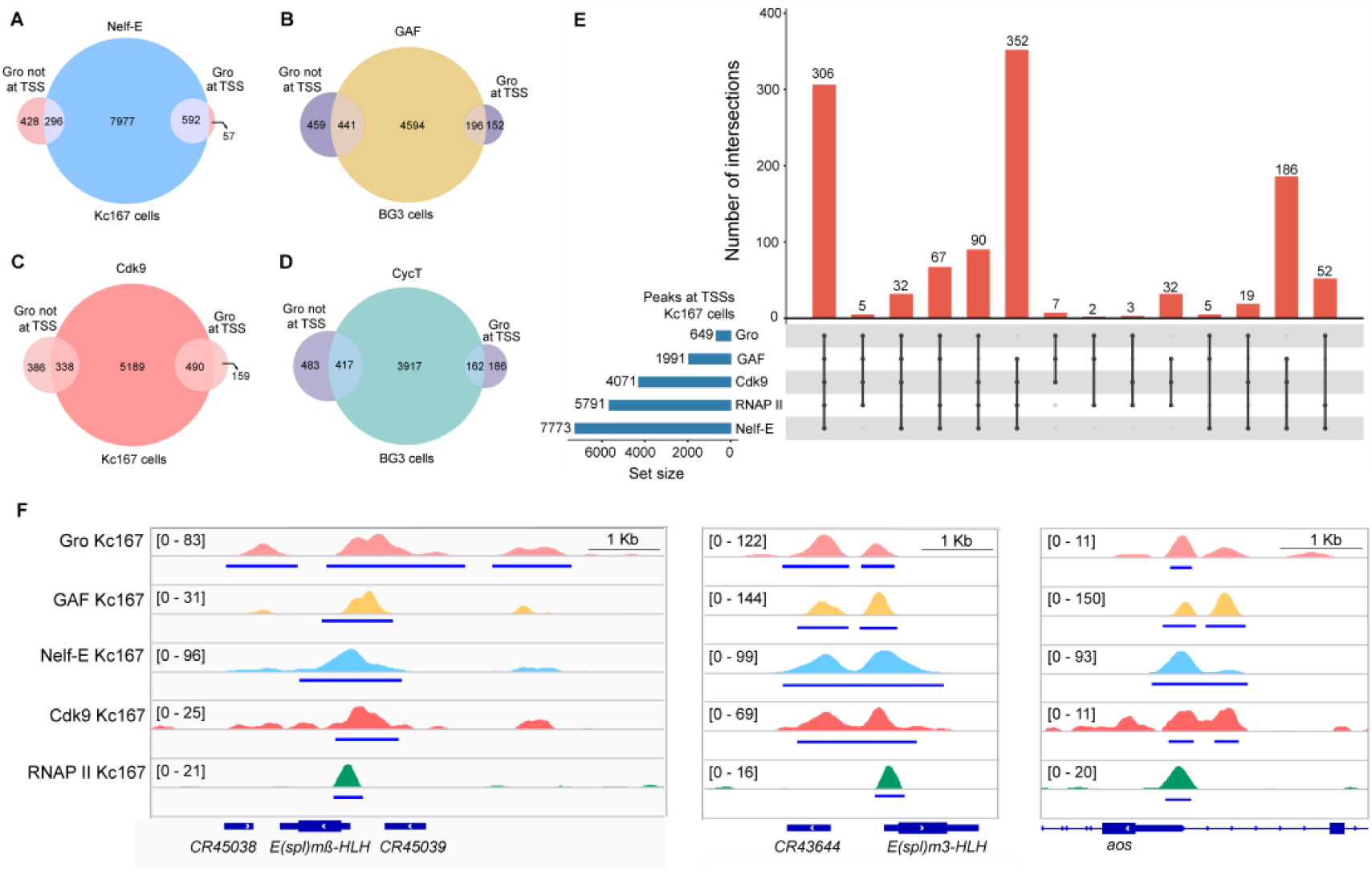
Gro co-occupies genomic locations with Nelf-E, GAF, Cdk9, CycT and RNAP II. A) Venn diagram showing the overlap between Gro and Nelf-E peaks located at GAF cdk9 and at all other sites (not at TSSs) in Kc167 cells. B) Venn diagram illustrating the overlap between Gro and GAF peaks located at TSSs and at all other sites (not at TSSs) in BG3 cells. C) Venn diagram showing the overlap between Gro and Cdk9 peaks located at TSSs and at all other sites (not at TSSs) in Kc167 cells. D) Venn diagram illustrating the overlap between Gro and CycT peaks at, and not at, the TSSs in BG3 cells. E) UpSet plot showing the intersections among peaks at TSSs of the following data sets: Gro, Nelf-E, Cdk9, GAF, and total RNAP II (Rpb3) The connected lines in the lower panel of the plot and the upper bar graph (in red) represent respectively the sets of peaks included in each intersection and its corresponding number of overlapping peaks. The horizontal left bar graph represents the total number of peaks in each data set. F) Genome browser views for Gro, GAF, Nelf-E, Cdk9, and total RNAP II (Rpb3) at *E(spl)mβ-HLH*, *E(spl)m3-HLH*, and *argos* (*aos*).

GAF is also linked to promoter proximal pausing at many genes in *Drosophila* and acts to promote pausing in part by recruitment of NELF (Li *et al*., 2013). Using GAF peaks derived from ChIP-seq data generated as part of the modENCODE project, we previously found that 82% of Gro peaks in Kc167 cells located at TSSs overlap with peaks of GAF (Kaul, Schuster and Jennings, 2014). In BG3 cells, we observe that 51% of Gro peaks overlap with GAF peaks, and this percentage is increased to 56% at TSSs (Figure 3B). These findings are consistent with the enrichment of GAF motifs at Gro peaks (Figure 1E) and indicate that Gro frequently occupies promoters bound by GAF, although this is not universal.

While GAF and NELF promote RNAP II pausing, release of the paused polymerase into productive elongation is driven by the P-TEFb complex, comprised of Cdk9 and Cyclin T (Peterlin and Price, 2006). Cdk9 is a kinase and acts to phosphorylate RNAP II, NELF and Spt5 on specific residues to release paused RNAP II into the elongation phase of transcription. One potential mechanism through which Gro could promote RNAP II pausing is by inhibiting the recruitment of P-TEFb to target genes. In this model, we would expect to see Cdk9 and Cyclin T peaks excluded from TSSs that bind Gro. We examined the overlap of Gro peaks with the Cdk9 ChIP-seq peaks in Kc167 cells (Nazer *et al*., 2018) and found 75.5 % of Gro peaks overlapped with Cdk9 peaks at TSSs (Figure 3C) and 46.6 % of Gro peaks overlapped with Cyclin T (Schaaf *et al*., 2013) at TSSs in BG3 cells (Figure 3D). Thus, Gro binding is frequently observed at promoters where P-TEFb components are also detected, arguing against a simple model in which Gro represses transcription by excluding P-TEFb from paused promoters.

To visualise the profile of pausing factors recruited with Gro at TSSs, we generated an Upset plot of overlapping peaks using published data from Kc167 cells for Nelf-E, Cdk9, GAF and RNAP II (Khan and Mathelier, 2017). Most frequently, Gro peaks overlapped with peaks of all these factors (Figure 3E; 306 of 649 Gro peaks). This combination of peaks is observed at three genes that are established Gro targets; *E(spl)mβ-HLH, E(spl)m3-HLH* and *aos* (Figure 3F) (Housden, Terriente-Felix and Bray, 2014; Kaul, Schuster and Jennings, 2014). The second most frequent combination observed included Nelf-E, Cdk9 and RNAP II peaks, but not peaks of GAF (Figure 2E; 90 of 649 Gro peaks).

Taken together, these results indicate that Gro frequently occupies promoters that are also bound by factors previously shown to regulate RNAP II pausing and pause release. The overlap of Gro with Nelf-E and GAF is consistent with our previous observations that Gro can influence promoter proximal pausing (Kaul, Schuster and Jennings, 2014). However, peaks of Cdk9 and Cyclin T are also found to overlap Gro peaks at TSSs of known Gro target genes, arguing against a model in which Gro promotes RNAP II pausing solely by preventing P-TEFb recruitment. Instead, these observations suggest that Gro functions within the canonical promoter proximal pausing environment and may modulate progression through the early elongation checkpoint rather than blocking assembly of the pause release machinery.

### Establishment of an assay to test for genetic interactions with *gro* during wing development

Sensitised genetic backgrounds in *Drosophila* are widely used to detect functional interactions through modification of reproducible developmental phenotypes (Johnston, 2002). To test whether the association of Gro with RNAP II pausing regulators observed in cultured cells reflects a functional relationship in vivo, we used a wing-specific knock-down of *gro* to assess genetic interactions with pausing factors using the *GAL4/UAS-RNAi* system (Heigwer, Port and Boutros, 2018).

We used a combination of *nub-GAL4* and *UAS-RNAi-gro* to knock down *gro* expression in the developing wing (Figure 4). *nub-GAL4* is expressed in the developing wing pouch region of the wing imaginal disc (Csordás, Grawe and Uhlirova, 2020). Flies carrying a single copy of each of *nub-GAL4* and *UAS-RNAi-gro* exhibited mild but reproducible defects in wing patterning (Figure 4B). We observed ectopic venation above the L2 vein, frequent loss of the anterior cross-vein (ACV), and weak spreading of longitudinal veins (Figure 4B). Increasing the dosage to two copies of *nub-GAL4* and two copies of *UAS-RNAi-gro* resulted in more penetrant and severe phenotypes (Figure 4C), including thickened longitudinal veins extending further at the wing margin, additional ectopic veins in the posterior wing, and complete loss of the ACV.

**Figure 4:**
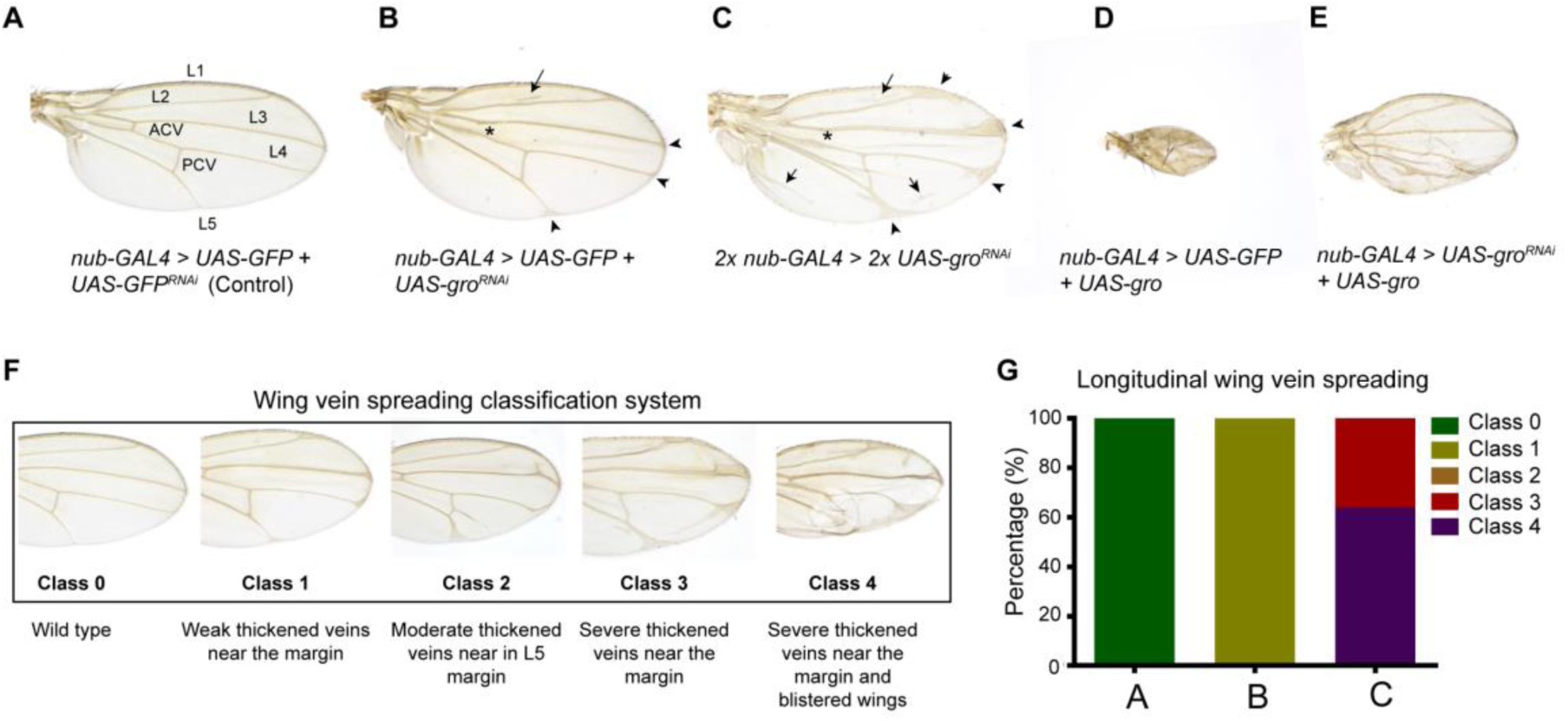
Characterization of *gro* RNAi knockdown during wing development. Representative examples of *Drosophila* adult female wings raised at 25^ο^C of the indicated genotypes. (A) Control wing showing key structures: anterior cross vein (ACV), posterior cross vein (PCV), and longitudinal veins L1 to L5. (B) Heterozygous *gro* RNAi knockdown showing ectopic venation above L2 (arrow), lack of ACV (asterisk) and weak longitudinal vein spreading (arrowheads). (C) Homozygous *gro* RNAi knockdown showing ectopic venation above L2 (arrow), increase of ectopic venation in the posterior half of the wing (arrows), loss of the ACV (asterisk), and severe longitudinal vein spreading (arrowheads). (D) *gro* overexpression showing a small and blistered wing. (E) Simultaneous *gro* knockdown and overexpression showing partial rescue of the wing size and wing vein pattern. (F) Adult wing images that are representative for each of the classes in the classification system and a brief description of their phenotypes. The classification system is based on the severity of the longitudinal vein spreading. There are four classes, which represent increasing degrees of severity (Class 0, 1, 2, 3 and 4). (G) Graph showing the frequency of each class observed for the following phenotypes: control [A], single *gro* knockdown [B], and double *gro* knockdown [C]. At least 70 wings of each genotype were scored for quantification in panel, except for c for which only 25 wings were scored due to the difficulty of obtaining viable flies.

The vein thickening and ectopic vein phenotypes are consistent with gro loss of function clones in the wing (Celis and Ruiz-Gómez, 1995) and the established model that *gro* acts as an effector of the Notch signalling pathway to repress vein cell fate during wing development (Hasson and Paroush, 2006). The reproducible loss of the ACV further indicates a requirement for Gro during later stages of wing patterning, including pupal development when cross-vein formation occurs (Blair, 2007).

To test the ability of the *UAS-RNAi-gro* line to knock down *gro* expression, we tested its ability to rescue the phenotype produced by overexpression of Gro via *UAS-gro*. Flies carrying *nub-GAL4* and *UAS-gro* exhibited small, blistered wings (Figure 4D). This phenotype was largely rescued by a single copy of *UAS-RNAi-gro*, confirming that the *UAS-RNAi-gro* transgene targets *gro* mRNA (Figure 4E). After the initial characterisation the *nub-GAL4; UAS-RNAi-gro* phenotypes, we established a system to classify the severity of the wing phenotypes observed to facilitate our analysis of genetic interactions using GAL4/UAS-RNAi lines (Figure 4F, G). This sensitised background provided a platform to test predictions of the early elongation checkpoint model in vivo.

### Knock-down of genes encoding factors that promote RNAP II pausing enhance *gro* phenotypes during wing development

Having established a genetic assay to test for interactions with *gro,* we next tested whether reducing the function of genes encoding regulators of RNAP II pausing modifies the gro phenotype in a manner predicted by the pausing model. If Gro attenuates transcription by promoting or stabilising promoter proximal pausing, then partial reduction of pausing factors would be expected to enhance the *gro* knock-down phenotype.

We have observed that peaks of Gro recruitment to the genome frequently overlap peaks of GAF recruitment in both Kc167 and BG3 cells (Figure 3B)(Kaul, Schuster and Jennings, 2014), but it is not known if this co-localization has functional significance *in vivo*. Expression of *Trl* (which encodes GAF) was reduced alone or in combination with *gro* using *nub-GAL4* (Figure 5A). Consistent with previously published work (Blanch, Pineyro and Bernues, 2015), knock-down of *Trl* alone resulted in reduced wing blade size but had minimal effects on vein patterning. In contrast, simultaneous knock-down of *gro* and *Trl* resulted in enhanced vein patterning defects relative to *gro* knock-down alone (Figure 5A), including increased longitudinal vein spreading (Figure 5B), ectopic venation and loss of the ACV (Figure 5C). The severity of these phenotypes resembled that observed upon increasing gro knock-down dosage. These findings are consistent with a functional interaction between Gro and GAF during wing vein patterning.

**Figure 5:**
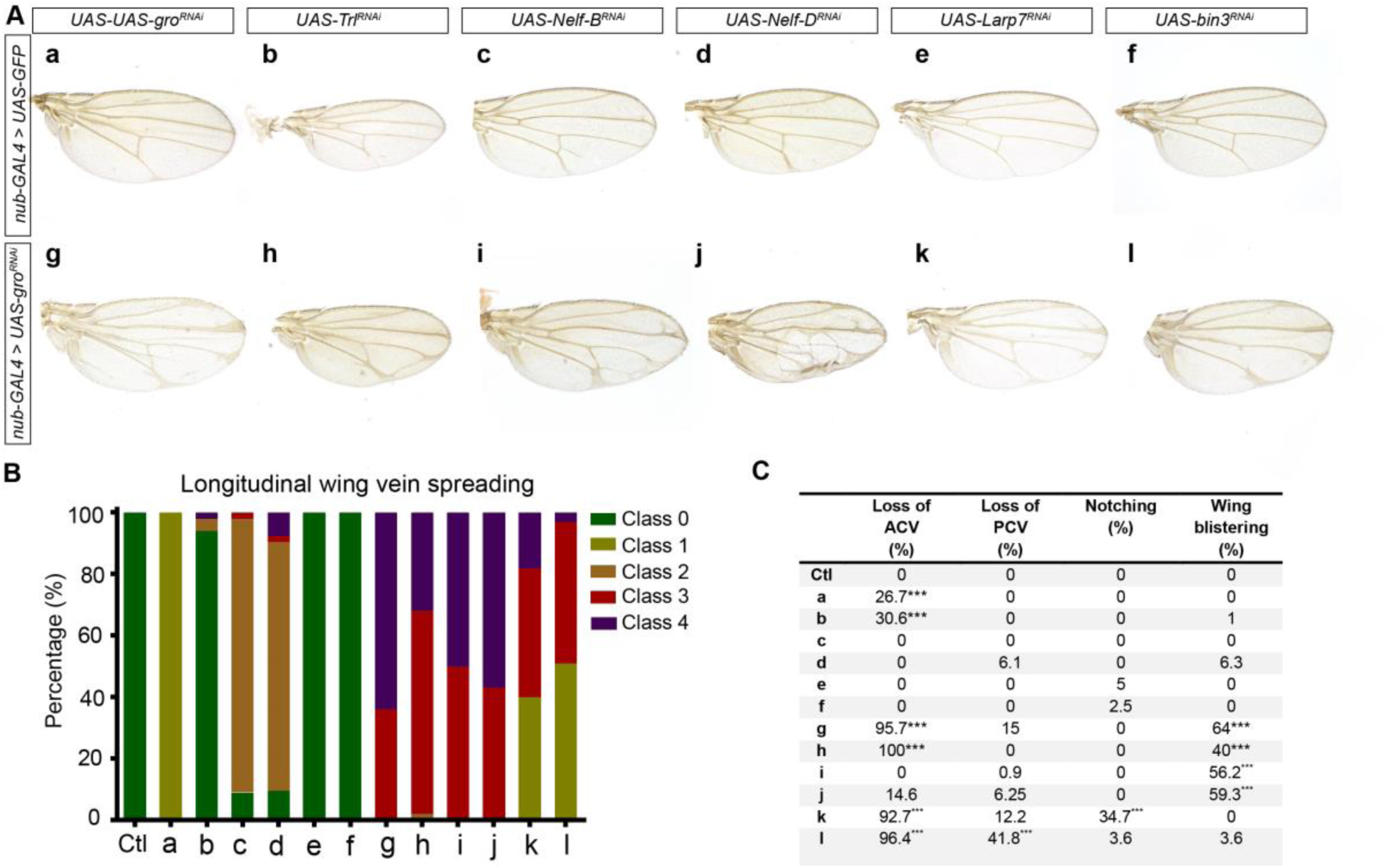
*gro* genetically interacts with *NELF*, *Trl*, *Larp7*, and *bin3* in the developing wing. (A) Representative examples of *Drosophila* adult female wings raised at 25^ο^C expressing the indicated transgenes. (B) Graph showing the frequency of each class (as defined in Figure 4) observed for the phenotypes shown in A. (C) Table showing the quantification of four wing features [total or partial loss of anterior cross vein (ACV), total or partial loss of posterior cross vein (PCV) notching, and wing blistering] for the phenotypes shown in A. Fisher’s exact tests were used for pairwise statistical comparisons. For each feature, the single knockdowns were compared to the control, and double knockdowns were compared to the single knockdowns of the genes in the double. ***p<0.0001. At least 70 wings of each genotype were scored for quantification in B and C.

We tested the knock-down of all 4 subunits of *Drosophila* NELF (A, B, D and E) for interactions with *nub-GAL4:UAS-RNAi-gro.* In this assay, the knock-down of either *Nelf-B* or *Nelf-D* alone resulted in weak but consistent phenotypes, and both significantly enhanced the *nub-GAL4:UAS-RNAi-gro* phenotype (Figure 5A, B, C). The knock-down of either *Nelf-A* or *Nelf-E* did not result in a visible phenotype or strong modification of the *nub-GAL4:UAS-RNAi-gro* phenotype (Supp Figure 2). However, we observed a weak increase in vein thickening in the double knock-down *gro* and *Nelf-A.* These differential effects may reflect variation in RNAi efficiency or distinct subunit requirements during wing development.

The 7SK small non-coding RNA (snRNA) inhibits RNAP II transcription by binding P-TEFb and recruiting an RNA binding protein HEXIM (HEXIM1 or HEXIM2 in mammals) to block Cdk9 activity (Peterlin, Brogie and Price, 2012). The association of P-TEFb with the 7SK snRNP complex is reversible and regulation of this association is a key mechanism to control P-TEFb activity in cells. We tested the genes encoding La-related protein 7 (Larp7, a component of the 7SKsnRNA complex), Bicoid-interacting protein 3 (Bin3, an RNA methyltransferase that adds a 5’-methyl cap to the product of snRNA:7SK preventing its degradation) and HEXIM for genetic interactions with *gro as UAS-RNAi lines* were available for these genes.

Knock-down of either *Larp7* or *bin3* alone did not lead to any detectable phenotypes in the wing. However, when either of these genes were knocked-down in combination with *gro,* there was a significant enhancement of the phenotypes observed in *nub-GAL4>UAS-RNAi-gro* flies (Figure 5). Knock-down of HEXIM using *nub-GAL4>UAS-RNAi-HEXIM* was lethal, and no flies were recovered to analyse.

Taken together, reduction of multiple regulators of RNAP II pausing enhances the *gro* knock-down phenotype during wing development. These genetic interactions are consistent with the hypothesis that Gro attenuates transcription in vivo in the context of the promoter proximal pausing machinery. While these data do not establish a direct molecular interaction, they support a model in which Gro acts with regulators of the early elongation checkpoint to influence transcription during wing development.

## DISCUSSION

Gro/TLE proteins have long been recognised as co-repressors for many different transcription factors that direct developmental processes. Here, we show that although Gro is recruited to genes in a largely cell-type specific manner, Gro target sites share common architectural features across cell types, including focal binding within accessible, enhancer-associated chromatin, providing insight into its mechanism of action.

The finding that Gro is predominantly recruited to open chromatin across different cell types is consistent with the long-established model that Gro is recruited by sequence-specific DNA-binding transcription factors. Most transcription factors known to recruit Gro [including the E(spl)-bHLHs, Capicua, and Brinker, involved with vein patterning] are not considered to be pioneer factors as they are unable to bind sequences in highly compacted closed chromatin (Hasson and Paroush, 2006; Balsalobre and Drouin, 2022). The association of Gro with open chromatin is also consistent with the rapid reversibility of Gro-mediated repression. For example, Gro forms part of the Su(H)/H complex that represses Notch targets in the absence of Notch signalling in *Drosophila* (Barolo *et al*., 2002; Nagel *et al*., 2005). Repression by this complex is rapidly reversed (within minutes) when Notch signalling is activated to enable Notch target gene expression (Krejci *et al*., 2009). Similarly, during cell cycle progression, Gro represses genes such as *e2f1* during S phase, and this repression is relieved by Cdk1 activity to permit timely progression into G2 and mitosis (Bar-Cohen *et al*., 2023). These examples illustrate that Gro-mediated repression operates in dynamic regulatory contexts where transcription must be tightly controlled yet rapidly reversible. Such rapid reversibility is difficult to reconcile with models invoking large-scale chromatin compaction but is compatible with regulatory mechanisms that act at the level of transcriptional elongation. The promotion of RNAP II pausing represents one such reversible mechanism, as exemplified at heat shock genes (O’Brien and Lis, 1991; Himanen and Sistonen, 2019).

Our analysis of ChIP-seq data reveals that Gro peaks frequently overlap peaks of factors implicated in promoter proximal pausing, including NELF and GAF. However, the presence of Gro does not exclude RNAP II or P-TEFb recruitment (Figure 3). For example, we observe overlapping peaks of Cdk9 and Gro binding around the TSS of *E(spl)HLH-mβ* in Kc167 cells (Figure 3F). Depletion of Gro has been shown to induce the release of paused RNAP II and increased transcription of *E(spl)HLH-mβ* in this cell line (Kaul, Schuster and Jennings, 2014), revealing that Cdk9 is not fully active when Gro is functioning as co-repressor at this locus. Together, these observations are consistent with a model in which Gro influences the early elongation checkpoint rather than preventing recruitment of elongation factors.

Studies in homogeneous cell populations have provided valuable insight into the molecular basis and prevalence of RNAP II pausing, but they do not establish whether modulation of pausing plays functionally significant roles during animal development. Genetic interaction analysis provides a complementary in vivo approach to assess functional relationships between gene products and has been central to defining developmental pathways (Pérez-Pérez, Candela and Micol, 2009). Null or strong loss-of-function mutations in core elongation factors are frequently lethal, limiting their usefulness for developmental genetic analysis. We therefore used the GAL4:UAS-RNAi system to reduce, rather than eliminate, gene expression and examine modification of a sensitised *gro* wing phenotype.

The single knock-down of *gro* (*nub-GAL4/+:UAS-RNAi-gro/+)* resulted in a mild but reproducible wing phenotype that could be modified by the concurrent knock-down of other genes by *UAS-RNAi* (Figures 4, 5). In several cases, the severity of the double knock-down phenotypes exceeded that expected from simple addition of the individual single knock-down effects, indicating a non-additive, synergistic interaction. Such non-additive enhancement is consistent with partial impairment of a shared biological process (Pérez-Pérez, Candela and Micol, 2009). In the wing, Gro functions to repress vein-promoting genes in intervein cells; reduction of Gro levels therefore weakens repression in this domain. Concurrent reduction of factors operating within the same repressive mechanism would be expected to further limit residual Gro function to further compromise repression and produce vein phenotypes exceeding additive effects.

Depletion of Gro by RNAi in the developing wing predominantly affects vein development, consistent with Gro’s role as a key effector of Notch signalling during vein patterning (Hasson and Paroush, 2006). The knockdown of individual factors known to promote pausing led to indiscernible or weak phenotypes in the wing, but concurrent knockdown of these factors with *gro*, greatly enhanced the *gro* vein phenotypes (Figure 5). These genetic interactions are consistent with the interpretation that Gro and pausing factors contribute to a shared repressive mechanism operating during wing vein patterning.

Co-localization of ChIP peaks indicates that proteins are recruited to overlapping genomic regions and have the potential to function together, but it does not establish functional interdependence. Synergistic genetic interactions provide evidence that gene products function in the same pathways but do not necessarily reflect a direct physical interaction. Here we present evidence that Gro recruitment to the genome co-localizes with that of pausing factors and that concurrent knockdown of *gro* and pausing factors results in synergistic phenotypes. Drawing from these two lines of evidence, we can deduce that Gro functions with factors that promote RNAP II pausing during wing patterning either via direct physical interactions or indirectly though yet unidentified intermediaries.

We propose that Gro attenuates transcription at the P-TEFb–dependent early elongation checkpoint. In this model, sequence-specific transcription factors recruit Gro to accessible regulatory regions. Gro acts within the context of promoter proximal pausing machinery, potentially reinforcing pausing or limiting productive P-TEFb activity, thereby restricting transition of RNAP II into productive elongation. This mechanism would permit rapid reversibility of repression upon receipt of activating signals that stimulate elongation. Such a model reconciles the focal, enhancer-associated binding of Gro with its established role as a transcriptional repressor and is inconsistent with historical models proposing extensive spreading of Gro across chromatin.

While it has become well established that many genes in developmental and stimulus-responsive pathways exhibit RNAP II pausing, insight into how RNAP II pausing is established in a gene-specific manner remains scarce (Gaertner and Zeitlinger, 2014; Core and Adelman, 2019; Dollinger and Gilmour, 2021). Gro is recruited by many transcription factors that are expressed in response to developmental and stimulus-responsive pathways. Our findings support a model in which Gro contributes to transcriptional attenuation by intersecting with pausing machinery, providing a mechanism through which diverse transcription factors may influence elongation control. Whether this represents a predominant mode of Gro/TLE function in other developmental contexts or in mammalian systems remains to be determined.

## MATERIALS AND METHODS

### Fly stocks and maintenance

The conditional knockdown of specific genes was achieved using the GAL4/UAS system (Brand and Perrimon, 1993). The following GAL4 and UAS-RNAi and UAS stocks were obtained from Bloomington Drosophila Stock Centre (BDSC); stock numbers indicated in brackets: *nub-GAL4* (86108), *UAS-Nelf-D^RNAi^* (38934), *UAS-Nelf-B^RNAi^* (34847), *UAS-Nelf-A^RNAi^* (32897), *UAS-Nelf-E^RNAi^* (32835), *UAS-Bin3^RNAi^* (50634), and *UAS-GFP* (4775). UAS-RNAi stocks used were also obtained from Vienna Drosophila Resource Center (VDRC; www.vdrc.at; stock numbers indicated in brackets): *UAS-gro^RNAi^* (6316), *UAS-Trl ^RNAi^*(41095), *UAS-Larp7 ^RNAi^* (31635). *UAS-gro* (Orian *et al*., 2007) was a gift from Ze’ev Paroush.

All flies were raised on standard cornmeal food and maintained either at 25^ο^C or at 18^ο^C in an incubator. Crosses were set up at 25^ο^C on standard cornmeal media with addition of dry yeast.

### Selection of the *UAS-RNAi gro* line

We tested three *Drosophila* transgenic RNAi lines against *gro* from the Vienna Drosophila Resource Center (VDRC) (fly stocks 6316 and 6315 from the GD RNAi collection, and 110546 from the KK RNAi collection), and one from the Bloomington Drosophila Stock Center (fly stock 91407). We excluded the fly line from the KK RNAi collection (110546) as it expresses ectopic *tiptop* (Vissers et al., 2016), as well as the one from Bloomington as it gave no phenotype when driven by *nub-GAL4.* The stocks from the VDRC GD RNAi collection (6316 and 6315) gave a very similar phenotype to each other when driven by *nub-GAL4*. However, the 6315 line proved difficult to maintain as an expanded stock so we used 6316 in our experiments, which we refer to as *UAS-gro^RNAi^.* We included *UAS-GFP* in our crosses to maintain a constant number of UAS elements in the single and double knockdowns to ameliorate any effects due changes in the ratio of GAL4 to UAS binding sites across the genome.

### Analysis of ChIP-seq data

Sequences were aligned to the dm6.26 reference genome using bowtie2 v2.3.3 (Langmead and Salzberg, 2012) with the default parameters, specifying the options “--no-unal” and “-local” to remove unmapped reads and trim adapters. Multimapping reads were then removed using the “view” program of SAMtools v1.7 (Li *et al*., 2009) with the parameter “-q 20”. The program “markdup” was also used to remove PCR duplicates from mapped reads, with the parameter “-r”.

For the analysis of Gro ChIP-seq data in BG3, two biological replicates were used to obtain a high confidence set of peaks. Peaks were called using the MACS2 v2.1.1.2 software providing input and immunoprecipitated samples simultaneously with default parameters. Peaks present in both biological replicates (with an FDR less than 10% in at least one of the samples and a p-value less than 0.001 in the other) were selected (Gaspar, 2018).

ChIP-seq peaks for the rest of the factors analysed in this work were retrieved from public databases (see Table 1). Peaks from the data bases that were aligned to the dm5 version of the *Drosophila* genome were converted to dm6.26 using the coordinate converter from Flybase (Larkin *et al*., 2020). Where ChIP-seq peaks from two replicates were provided, BEDTools intersect was used to obtain a single set of high confidence peaks using a minimum and reciprocal fraction of overlap of 5%.

**Table 1.**

| **Deposited data** |  | **GSE accession number** | **Reference** |
| --- | --- | --- | --- |
| Gro ChIP-seq | Kc167 | Array Express<br>E-MTAB-2316 | (Kaul et al., 2014) |
| Gro ChIP-seq | BG3 | Array Express<br>E-MTAB-12108 | (Bar-Cohen *et al.*, 2023) |
| Gro ChIP-seq | S2R+ | Array Express<br>E-MTAB-2316 | (Kaul et al., 2014) |
| GAF ChIP-seq | Kc167 | modEncode ID: 3245 | (Celniker *et al.*, 2009) |
| GAF Chip-Chip | BG3 | modEncode ID: 2651 | (Celniker *et al.*, 2009) |
| Nelf-E ChIP-seq | Kc167 | GSE116889 | (Nazer et al., 2018) |
| Cdk9 ChIP-seq | Kc167 | GSE116889 | (Nazer et al., 2018) |
| Pol II isoforms ChIP-seq | Kc167 | GSE116889 | (Nazer et al., 2018) |
| DNA-seq | BG3 | GSE41351 | (Lubelsky et al., 2014) |
| ATAC-seq | Kc167 | GSE122575 | (Porcelli *et al.*, 2019) |
| CycT Chip-Chip | BG3 | GSE42399 | (Schaaf et al., 2013) |

The Bioconductor library ChIPseeker v1.24.0 (Yu, Wang and He, 2015)was used to annotate the peaks by identifying nearby genes, and to visualize their genomic locations. Promoters were defined −250 to 250 bp from the TSSs.The breadth of Gro peaks in BG3 cells was analysed and plotted using the Bioconductor library ChIPpeakAnno v3.22.2 (Zhu *et al*., 2010). For optimal visualization, the histogram was divided into 500 bins and the two Gro peaks that were greater than 4000 bp were excluded from the histogram.

Open chromatin regions at Gro binding sites were analysed using a DNA-seq dataset in BG3 from modEncode (accession number: 5528), and an ATAC-seq dataset in Kc167 cells (Porcelli *et al*., 2019). DNA-seq and ATAC-seq signals at Gro binding sites were plotted in an average plot and a heatmap using computeMatrix and plotHeatmap from deepTools v3.4.3 (Ramírez *et al*., 2014).The plots were centered at the centre of Gro peaks and extended 2 Kb upstream and downstream of the reference point.

CentriMo v5.4.1 (Bailey and Machanick, 2012) software from the MEME suite was used to identify sequence motifs that were enriched in 500 bp sequences centred on the Gro binding peak summits in BG3 identified by MACS2. The Database for Annotation, Visualization, and Integrated Discovery (DAVID) v6.8 (Huang, Brad T. Sherman and Lempicki, 2009; Huang, Brad T Sherman and Lempicki, 2009) was used for the functional annotation of genes present at Gro binding sites or nearby identified by ChIPseeker.

The chromatin states files for the *Drosophila* genome in Kc167 and BG3 cells files were provided by Sarah Bray (Skalska *et al*., 2015).To analyse the chromatin states at Gro binding sites in those cell lines, BEDTools intersect was used with a minimum overlap of 50%. Percentage of Gro peaks within each chromatin state (or colour) were then plotted using GraphPad Prim v6.

### Co-localization study of ChIP-seq peaks

BEDTools intersect or Intervene (Khan and Mathelier, 2017) were used for the analysis of overlapping binding sites between Gro and the rest of the factors. A minimum and reciprocal fraction of 5% of overlap was required to consider it an overlap. The Venn diagrams shown in the figures were created using the matplotlib-venn package (Hunter, 2007). UpSet plots were generated directly from the Intervene module UpSet.

BigWig files were generated for the visualization of the genome tracks of ChIP-seq signal in the Integrate genomics viewer (Thorvaldsdóttir, Robinson and Mesirov, 2013). BigWig files were generated from BAM files using bamCoverage from deepTools v3.4.3 using a bin size of 10, and normalization by counts per million reads mapped (CPM). BigwigCompare was then used to normalize each file against its input using a bin size of 10 and operation subtract as parameters.

## ACKNOWLEDGEMENTS

We would like to thank Aamna Kaul for generating the Gro ChIP-seq data from BG3 cells and Christine Ashton for technical support with maintaining fly stocks and ordering. We thank Ze’ev Paroush, the Bloomington Drosophila Stock Center and the Vienna Drosophila Resource Center (VDRC) for providing fly stocks used in this study. The microscopy was performed with support from the Oxford Brookes University Bioimaging Centre. We acknowledge FlyBase and the modENCODE consortium for providing publicly available datasets. Many thanks to Alex Buffry, Sebastian Kittelmann, Gabrielle Mastrolonardo Raymond, Alara Erenel, Ze’ev Paroush and Kate Lines for thoughtful comments on the manuscript and Rob White and Saad Arif for a valuable discussion during MLMQ’s PhD thesis defence. Finally, we are particularly grateful to David Ish-Horowicz for comments on an early version of this manuscript. BHJ also wishes to acknowledge his mentorship over many years and the opportunity to begin work on Groucho in his laboratory. David passed away in 2024 and is greatly missed.

---

**Supp Figure 1:**
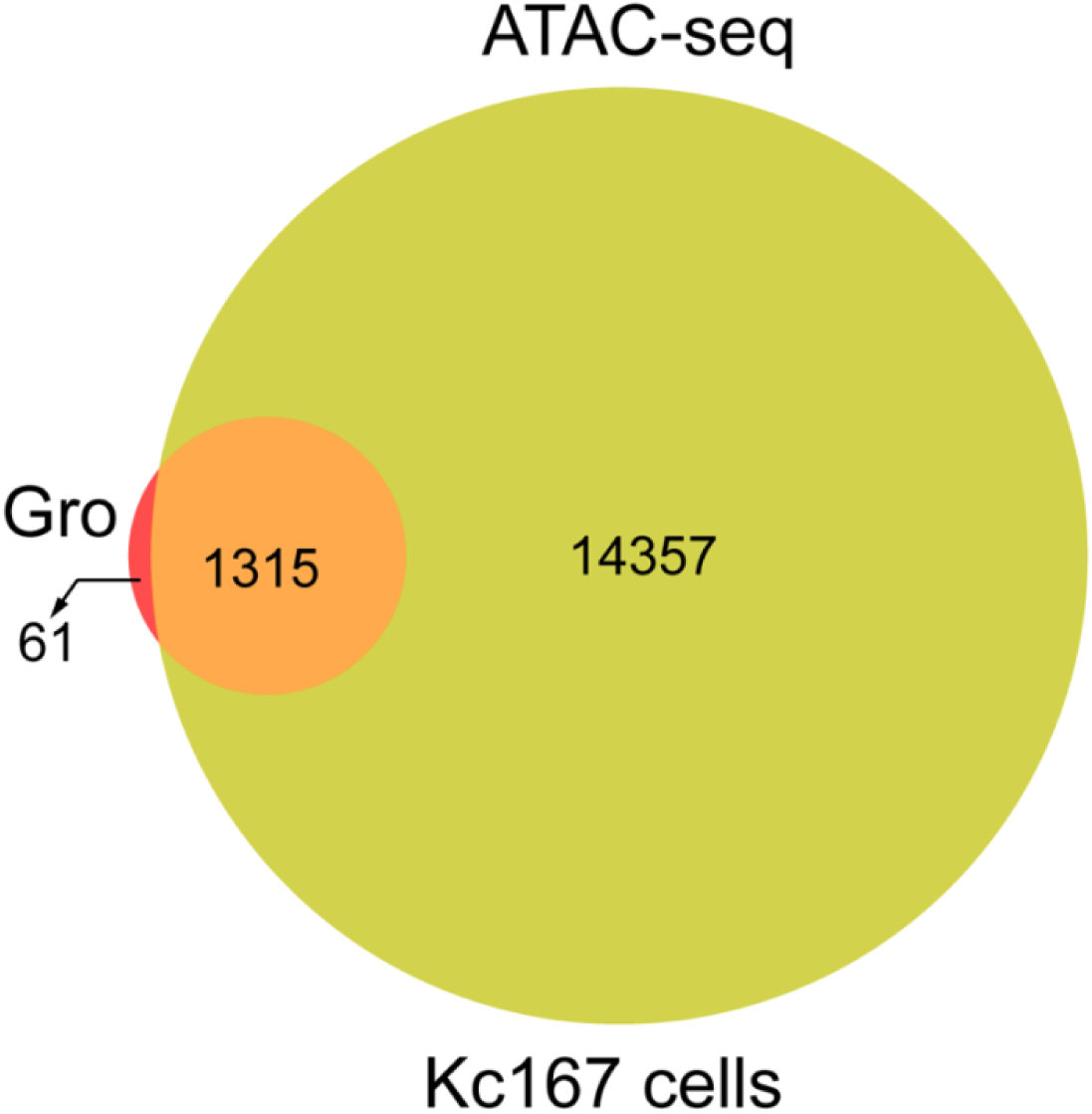
Gro peaks overlap ATAC-seq peaks in Kc167 cells. Venn diagram showing the overlap between Gro and ATAC-seq peaks in Kc167 cells.

**Supp Figure 2:**
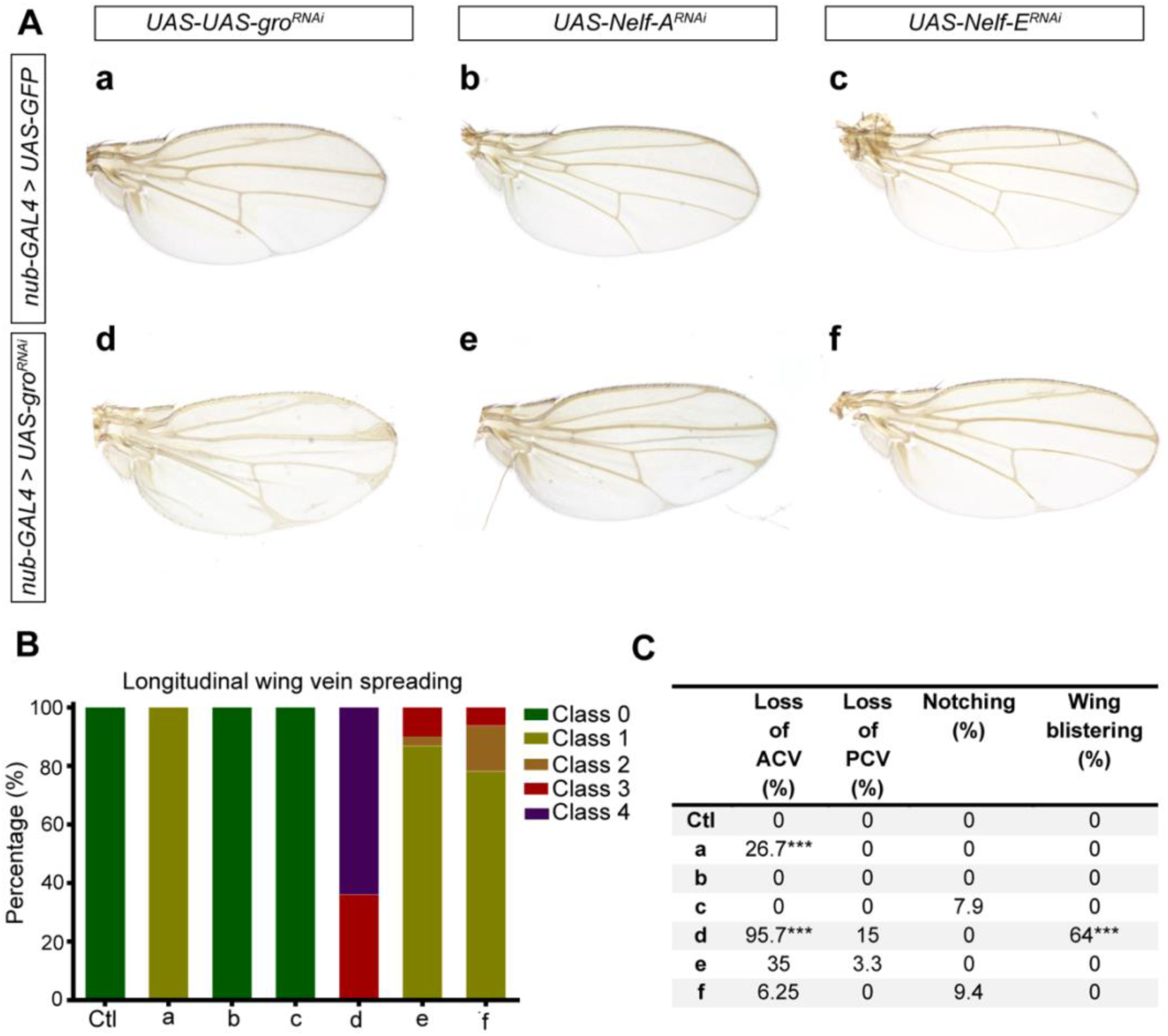
RNAi Knockdown *gro*, and *Nelf A/E* in the developing wing. (A) Representative examples of *Drosophila* adult female wings raised at 25^ο^C expressing the indicated transgenes. (B) Graph showing the frequency of each class (as defined in Figure 4) observed for the phenotypes shown in A. (C) Table showing the quantification of four wing features [total or partial loss of anterior cross vein (ACV), total or partial loss of posterior cross vein (PCV) notching, and wing blistering] for the phenotypes shown in A. Fisher’s exact tests were used for pairwise statistical comparisons. For each feature, the single knockdowns were compared to the control, and double knockdowns were compared to the single knockdowns of the genes in the double. ***p<0.0001. At least 30 wings of each genotype were scored for quantification in B and C.

